# Inhibition in motion: Test-retest reliability of inhibitory kinematics in a go/no-go mouse tracking task

**DOI:** 10.64898/2026.05.06.722889

**Authors:** Devu Mahesan, Komal Sharma, Maike Katharina Weinerth, Suman Dhaka, Marcus Meinzer, Rico Fischer

## Abstract

Response inhibition, the ability to suppress contextually inappropriate actions, is a cornerstone of cognitive control and is commonly assessed using paradigms such as the go/no-go task. However, traditional go/no-go paradigms rely on binary outcomes such as commission errors, which offer limited insight into the dynamic, graded behavioral adjustments underlying successful stopping. The present study developed a novel mouse-tracking go/no-go paradigm with a dynamic start to capture inhibitory processes during ongoing execution. Twenty-three healthy young adults completed the task in two sessions separated by approximately one week to evaluate the test-retest reliability of standard behavioral measures (error rates and reaction times), and three kinematic features: path length, mean velocity, and mean acceleration. Results revealed robust differences between go and no-go trials across all measures. Successful inhibition was characterized by significantly shorter path lengths and reduced mean velocity and acceleration compared to go trials. Critically, all measures demonstrated moderate-to-good test-retest reliability across sessions, with intraclass correlation coefficients ranging from .75 to .85 for go trials and from .59 to .83 for no-go trials. These findings establish construct validity and psychometric reliability of the current mouse-tracking go/no-go paradigm. The demonstrated stability of these measures provides the methodological foundation for their use in cross-sectional, longitudinal, and intervention research targeting inhibitory control.

## 1 Introduction

Humans rely on response inhibition to suppress contextually inappropriate actions that potentially interfere with goal-directed behavior in everyday life. In this context, the ability to withhold or override prepotent responses is considered a core component of cognitive control (Aron, 2011; Diamond, 2013; Duque et al., 2017; Mostofsky & Simmonds, 2008). One of the most frequently used paradigms to investigate inhibitory processes are go/no-go tasks that require participants to execute a response to a “go” stimulus and withhold responses to a “no-go” stimulus. Here, the relative frequency of go and no-go trials is used to create a strong prepotent response tendency, thereby isolating the inhibitory signal required to abort an action. Go/no-go tasks have been instrumental in neuroimaging research, suggesting that the right inferior frontal gyrus and anterior cingulate cortex are key nodes in the human inhibitory network (Aron et al., 2004, 2014; Chambers et al., 2009; Greenhouse et al., 2011). Apart from studying the cognitive and neural architecture of inhibitory processing, go/no-go paradigms have also been used extensively for studying inhibitory deficits in numerous neuropsychiatric conditions (Barkley, 1997; Nigg et al., 2005; Wright et al., 2014). Consequently, the go/no-go paradigm has become a cornerstone in experimental, neuroimaging, and clinical research.

Notably, to index inhibitory efficiency, traditional go/no-go paradigms frequently rely on binary behavioral outcomes, such as commission errors. Although the absence or presence of a response in no-go trials informs about successful or impaired inhibition, it provides only limited insight into the underlying cognitive processes. This limits the investigation of ongoing or corrective processes underlying inhibition (Leontyev et al., 2018; Leontyev & Yamauchi, 2019). Indeed, evidence from electromyography and neurophysiological studies demonstrates that inhibition is realized through graded adjustments in (motor) activity rather than discrete success or failure states (Allain et al., 2004; Cohen & van Gaal, 2014; Ficarella et al., 2019; Neely et al., 2017). Crucially, because binary outcomes only register whether a response was withheld or not, they primarily inform about failures of inhibition. Successful inhibition is treated as the absence of a response and remains largely uncharacterized. Continuous movement measures, by contrast, offer the possibility of examining how correct stopping unfolds, providing direct insight into the dynamics of successful inhibitory control.

This can be achieved by applying continuous measures of response times (RT) or mouse tracking to the study of inhibitory control. Specifically, based on suggestions that decisions and motor outputs develop jointly, investigation of continuous hand movement trajectories has been used to reveal the temporal dynamics (Scherbaum et al., 2010; Scherbaum & Dshemuchadse, 2020) in cued go/no-go (Benedetti et al., 2021; Bravi et al., 2022) and stop-signal paradigms (SST, Benedetti et al., 2020; Bravi et al., 2022; Leontyev & Yamauchi, 2019). For example, Benedetti et al. (2020) implemented a mouse-tracking procedure in a stop-signal and a cued go/no-go task, where advanced probabilistic cues signaled the likely trial type before the imperative stimulus appeared.

Movements were categorized from velocity profiles as one-shot (smooth, single-peaked) or non-one-shot (multi-peaked, corrected), providing an index of online corrections during execution. The authors found that inhibitory failures in the cued go/no-go task produced more frequent trajectory corrections than in the stop-signal task, suggesting that proactive control can disrupt the developing motor plan. Importantly, this work demonstrates that mouse-trajectory measures can capture aspects of inhibitory processing that binary outcomes miss, for example, the dynamic nature of inhibitory failure rather than merely recording a failure to stop. However, two important limitations of this work should be noted. First, the cued paradigm differs fundamentally from the standard go/no-go paradigm used in most experimental and clinical research, because advanced probabilistic information allows inhibitory preparation to begin before movement onset rather than being triggered by the stimulus during active movement. Second, because the analysis was restricted to classifying velocity profiles, the range of kinematic features that could be examined remained narrow. Mouse trajectories offer a richer array of scalar kinematic indices, such as acceleration profiles and total distance measures, which have been shown to be valuable for tracking individual and clinical variability (Leontyev et al., 2018; Leontyev & Yamauchi, 2019).

Finally, while prior work suggested that mouse tracking is well-suited to capture meaningful and dynamic aspects of inhibition, no previous study, to the best of our knowledge, has formally investigated the stability and reliability of go/no-go paradigms across a range of kinematic parameters. Previous studies have shown that traditional go/no-go measures, such as commission error rates and go RTs, exhibit acceptable test-retest reliability (Hedge et al., 2018; Langenecker et al., 2007; Wöstmann et al., 2013). However, no study, to the best of our knowledge, to date has examined whether kinematic features derived from continuous movement trajectories are similarly stable across sessions. This is not only relevant for cross-sectional biomarker research (e.g., studies investigating brain-behavior relationships), but also for longitudinal studies aimed at detecting subtle modulations of cognition by specific interventions (Meinzer et al., 2024). For example, neuromodulatory studies often report small effects on inhibition, and a recent meta-analysis of 45 transcranial direct current stimulation (tDCS) studies showed an overall small effect (g = .21) on inhibitory control and an even smaller effect for go/no-go paradigms (g = .10; Schroeder et al., 2020). Here, variability in outcomes has been attributed not only to stimulation parameters but also to interactions with task requirements and psychometric properties, including stability across sessions and reliability within individuals.

These current knowledge gaps were addressed in the present study: The first objective was to develop a mouse tracking go/no-go paradigm with a dynamic start, to ensure that inhibitory processes occur during active movement instead of being resolved before movement initiation. Such a design not only allowed us to measure RTs in no-go trials but also to investigate a range of dynamic kinematic features, such as mean velocity, mean acceleration, and path length, that serve as established summary indices of movement dynamics in mouse-tracking research (Kieslich et al., 2019; Wulff et al., 2025). Importantly, examining the temporal dynamics and kinematic features of no-go trial processing can provide the opportunity for a potential decomposition of the processes underlying successful inhibition (Sternberg, 1969). Our second objective was to determine test-retest reliability of these measures. For this purpose, participants completed the task twice across a one-week interval, to evaluate whether outcomes were stable and reliable across individuals across sessions.

## 2 Methods

### 2.1 Participants

A total of 26 participants took part in the study (15 female, 1 diverse; mean age: 22.9 years; range: 18 - 33 years). The sample consisted mainly of students from the University of Greifswald (*n* = 25), while one participant reported being employed. Each participant completed the study in two experimental sessions, separated by a mean test-retest interval of 9.6 days (range: 7 - 16 days). All reported normal or corrected-to-normal vision, and no neurological or psychiatric history. Right-handedness was verified prior to the first experimental session using the German version of the Edinburgh Handedness Inventory (Oldfield, 1971). All participants provided informed consent before participation and completed a brief demographic questionnaire. The study was conducted in accordance with the Declaration of Helsinki and the ethical guidelines of the German Psychological Society. Participants were compensated with either course credit or €20.

### 2.2 Apparatus and stimuli

The experiment was conducted on a Lenovo ThinkBook 14-IIL notebook PC, equipped with a 14-inch display (1920 × 1200 px, 60 Hz refresh rate), and was programmed in Psychopy Builder v2024.2.4. (Peirce, 2007). Participants performed the task using a standard USB optical mouse; we used an extension cable to ensure that the device could be moved freely and without restriction. Mouse sensitivity was set to the default condition (50% velocity). To ensure consistent surface friction and smooth cursor control, the mouse was operated on a DIN A2 paper surface placed flat on the desk.

Each trial began with a start box (100 × 100 px), which was located at the bottom center of the screen (coordinates: x = 0, y = –400), while the response box of identical size, colored white, appeared in the upper-right quadrant (x = 200, y = 300). The stimuli, which consisted of the digits 1 to 9, were also presented in black font within the response box. The digits 2, 3, 4, 6, 7, and 8 served as the go stimuli, while 1 and 9 served as the no-go stimuli.

### 2.3 Experiment Procedure

Each trial began with a fixation cross displayed for 500 ms, and a blue start box was presented at the bottom center of the display. Participants were instructed to move the cursor into the start box and hold it there for 500 ms, thus ensuring movement initiation from a consistent spatial location. Afterwards, the box turned black, indicating the trial onset. In parallel, the white response box appeared in the upper-right corner of the screen. The stimulus was presented in the white response box only after the cursor crossed a vertical threshold located 100 pixels above the start position. Such a movement-contingent stimulus presentation allowed us to examine inhibition during active movement, rather than a decision made before movement initiation (Scherbaum & Kieslich, 2018). Participants had a maximum of 1500 ms to respond, followed by a 1000 ms inter-trial interval during which the cursor was not visible. The experimental procedure is shown in Figure 1.

**Figure 1.**
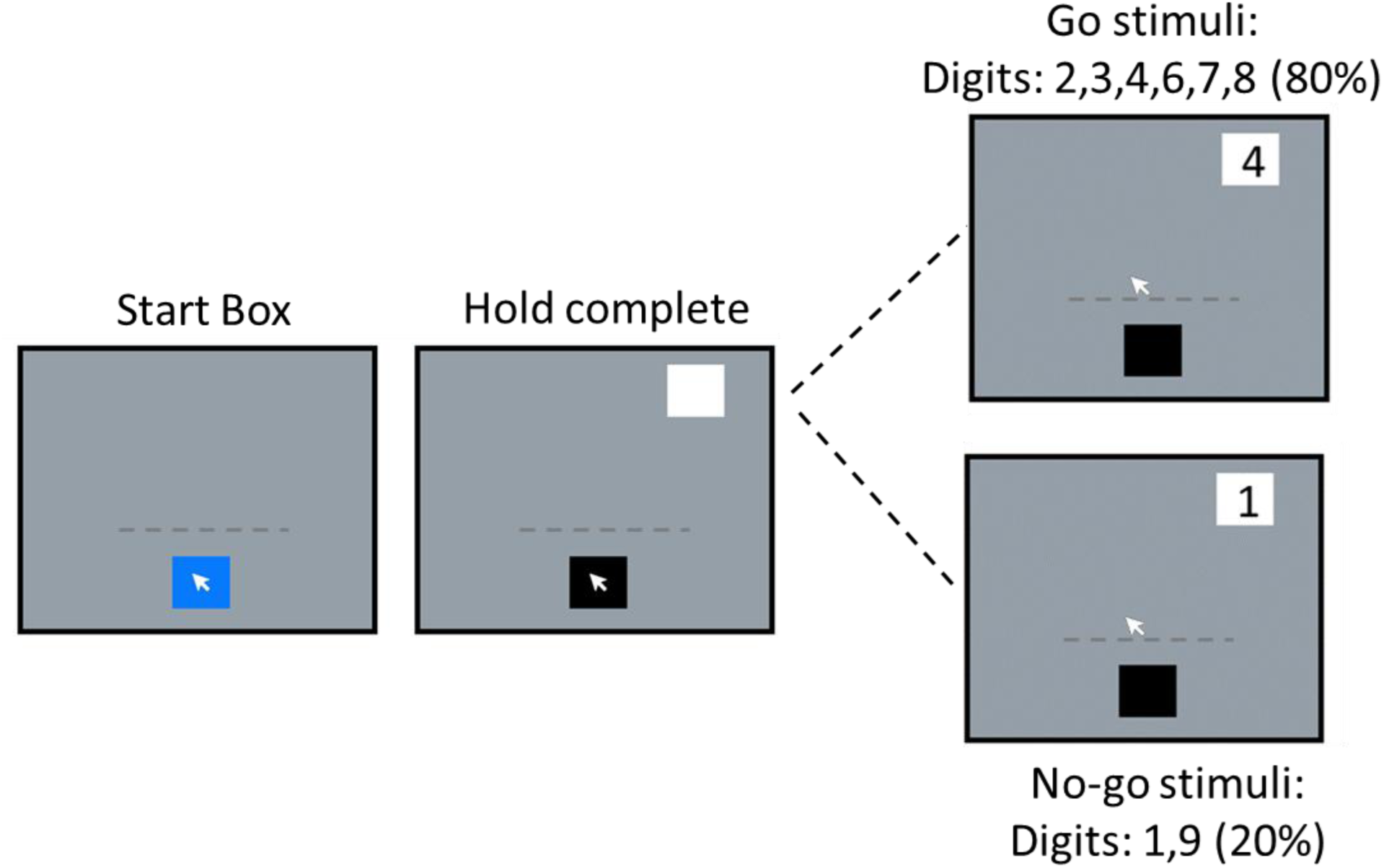
Schematic illustration of a trial sequence. *Note.* Each panel represents a successive stage of the trial. To begin, participants positioned the cursor within the start box and held it there to initiate the trial. Once the hold requirement was met, the start box turned black, indicating that movement could begin. As participants moved the cursor upward, the stimulus digit appeared in the response box only after the cursor crossed a predefined vertical threshold (represented here by the dashed horizontal line; note that this threshold was invisible to participants). On go trials, participants responded by clicking within the response box; on no-go trials, they were required to inhibit the ongoing movement and withhold the click.

Participants first completed two short practice blocks, one comprising only go trials and one comprising only no-go trials (18 trials each). The main experiment then consisted of repeated sequences of three blocks: one go-only block (18 trials) followed by two mixed go/no-go blocks (60 trials each; 80% go, 20% no-go). This sequence was then repeated four times. The session concluded with a final go-only block to achieve a symmetrical design, resulting in a total of five go-only blocks and eight mixed go/no-go blocks. Within each mixed block, go digits (2, 3, 4, 6, 7, 8) were each presented eight times, and no-go digits (1, 9) six times. Furthermore, trial order within the mixed block was pseudorandomized, where trial sequences were restricted to three possible transition types: go trial followed by a go trial, a go trial followed by a no-go trial, and a no-go trial followed by a go trial.

### 2.4 Design

We used a 2 (session: session 1, session 2) × 2 (trial-type: go, no-go) repeated measures within-subjects design. The present study reports error rates and RT as well as kinematic measures as dependent variables (please see below).

#### Error rates and RT

For go trials, a response was categorized as correct if the first left mouse click occurred within the response box. Go trials were scored as errors if: (a) the first left-click occurred outside the response box boundaries, (b) the click occurred with the right or middle button, or (c) no click was recorded within the deadline. Consequently, go RT is the time taken from stimulus onset until the correct click is made.

For no-go trials, accuracy was defined by successful motor inhibition and the absence of any mouse clicks. No-go trials were categorized as errors if: (a) any button press occurred during the trial duration, or (b) the participant failed to meet the stopping criteria. Correct stopping was computed using an approach informed by movement boundary-detection methods from human movement research (Wang & Welsh, 2024) and velocity threshold techniques previously applied in mouse tracking studies (Menceloglu et al., 2021; Wang et al., 2025). Specifically, we first identified the earliest point after the peak velocity at which the speed dropped below 10 px/s and remained below this level for 150 ms. We used a fixed velocity threshold (10 px/s) rather than a percentage-based rule (e.g., 3% of peak velocity) to ensure consistent stopping criteria across trials. To confirm that participants had fully stopped and to rule out any late corrective movements, we applied an additional verification step. To confirm sustained stopping, we required that at least 90% of velocity samples in the final 250 ms fall below the 10 px/s threshold. This added constraint helps address the known challenges in distinguishing true stopping (Wang & Welsh, 2024) from minor late adjustments or re-accelerations. Subsequently, the no-go RT, which is the inhibition latency, is the time taken from stimulus onset to the first moment of stopping. Given that stopping was operationalized as velocity dropping below 10 px/s for a continuous 150 ms window, the no-go RT corresponded to the start of this window.

#### Kinematic Measures and Preprocessing

For each trial, mouse trajectories were recorded as a sequence of x and y coordinates with their corresponding timestamps and processed in R. In order to preserve the dynamics of temporal data, all kinematic variables were calculated using raw-time analysis rather than time normalization (Hehman et al., 2015). This approach was adopted to preserve the temporal dynamics of movement, which is critical for accurately assessing the kinematic measures of response execution and inhibition.

To capture the dynamics of a response, we defined distinct epochs for data analysis of each trial type. For go trials, the epoch spanned from stimulus onset to the final click. For no-go trials, we utilized a stabilized epoch, i.e., from stimulus onset until 150 ms after the identified stop (or the end of recording). The additional 150 ms window ensured that kinematic measures contained the full braking phase, rather than stopping at the initial velocity threshold.

The kinematic dependent variables in the current experiment are path length, mean velocity, and mean acceleration. Path length is defined and computed as a measure of total distance traveled by the cursor during a trial. For this, the displacement between each pair of consecutive cursor samples was calculated, and these stepwise displacements were then summed across the entire trajectory. The resulting cumulative distance is considered the path length of a given trial. This measurement captures the overall spatial movement within a trial. Mean velocity was computed as the ratio of path length to the total movement duration. Movement duration was defined as the time elapsed between the first and last valid timestamp of the trial trajectory. Finally, acceleration was computed as the change in velocity between consecutive samples, divided by the corresponding time interval. To compute mean acceleration, absolute values were averaged using a time-weighted approach.

### 2.5 Data analyses

Data from practice blocks were not analyzed. Two participants did not attend the second session and were therefore excluded from all analyses. One additional participant was excluded due to low accuracy on go trials in the go/no-go block during session 1 (49%). Analyses were restricted to mixed go/no-go blocks to investigate inhibitory demands.

Error trials and outliers (2.1%) were removed prior to RT analyses. Outliers were identified within each participant × session × trial-type as values exceeding ± 2.5 standard deviations from the mean. A repeated measures ANOVA with the factors session (session 1, session 2) × trial-type (go, no-go) was conducted on each dependent variable. To evaluate test-retest reliability, we used the intraclass correlation coefficient based on a two-way mixed-effects model (ICC(3,1)), which is a well-established index appropriate for designs where the same participants complete identical measurements across two sessions (McGraw & Wong, 1996). For each dependent variable, session 1 and session 2 values were correlated separately for go and no-go trials. Following conventional benchmarks, ICCs are interpreted as follows: below .50 are considered poor, between .50 and .75 are considered moderate, between .75 and .90 are considered good, and above .90 are considered excellent (Koo & Li, 2016).

## 3 Results

### 3.1 Error rates and RT Error rates

Figure 2 illustrates the error rate as a function of session and trial-type, and Figure 3 represents the test-retest reliability between sessions for go and no-go trials. Participants made more errors in the no-go (12%) as compared to go trial (4%), *F*(1, 22) = 13.40, *p* = .001, *η_p_²* = .38. The main effect of session was not significant, *F*(1, 22) = 3.63, *p* = .070, *η_p_²* = .14, and the session × trial-type interaction was also not significant, *F*(1, 22) = 0.41, *p* = .527, *η_p_²* = .02.

**Figure 2.**
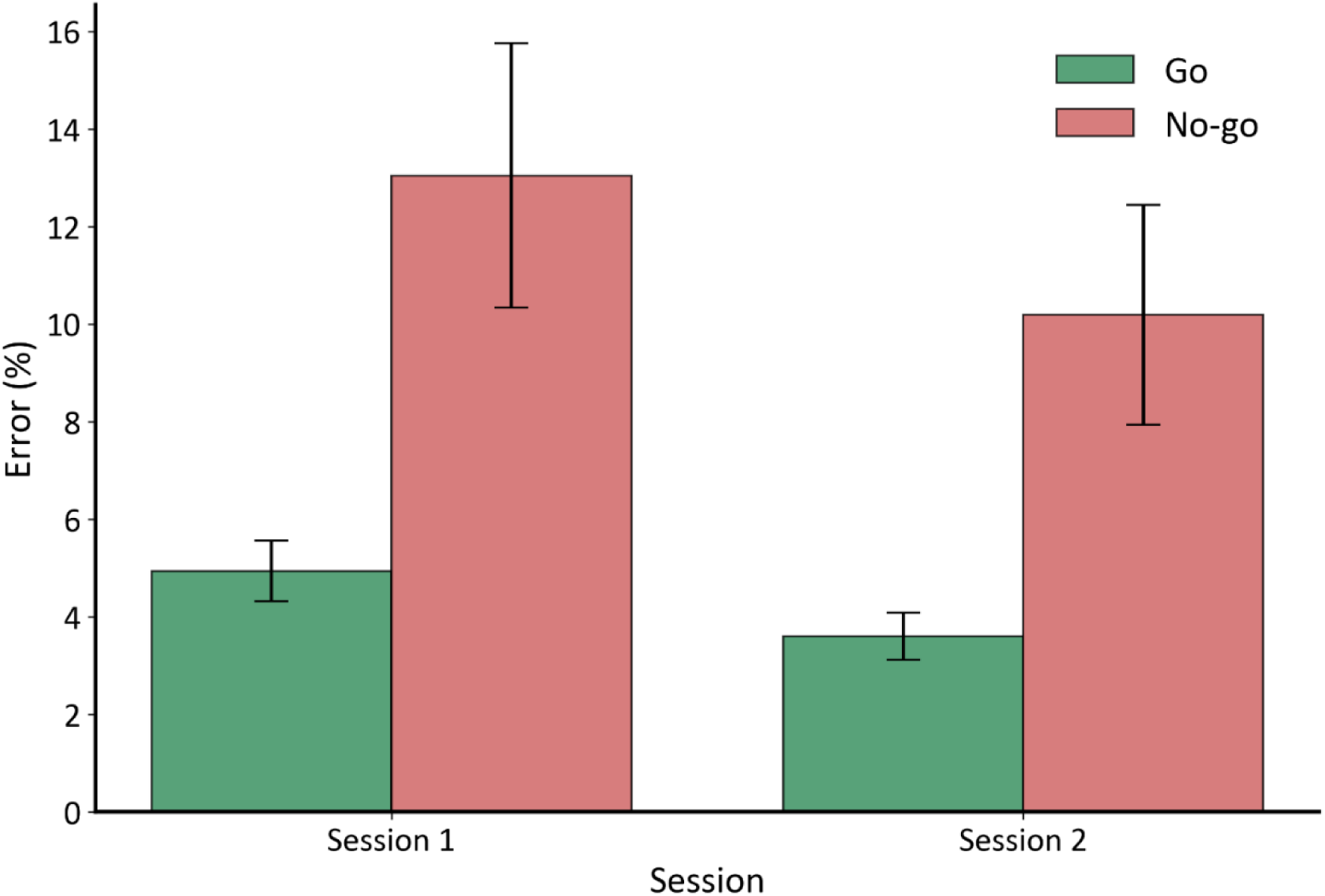
Error rate by session and trial-type. *Note*. Error rate (in %) as a function of session and trial-type. Error bars indicate the standard error of the mean (SEM) across participants.

**Figure 3.**
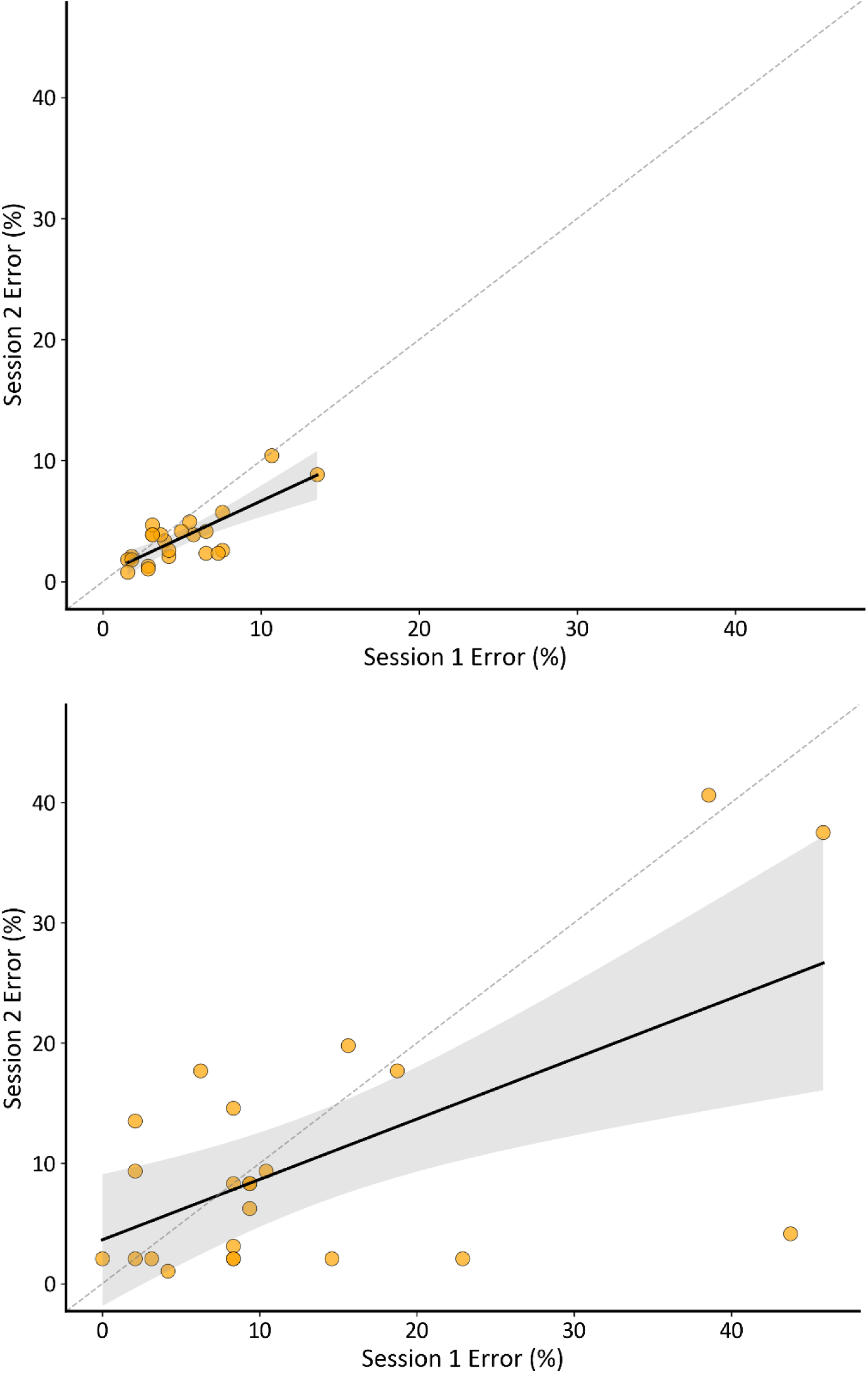
Test-retest reliability for go and no-go error rate between the sessions. *Note.* Scatter plot showing the relationship between session 1 and session 2 error rate (in %) for go (top) and no-go (bottom) trials. Each point represents one participant. The black line depicts the ordinary least squares regression of session 2 on session 1, and the shaded region indicates the 95% confidence interval of the mean regression estimate. The dashed diagonal line represents the line of identity (y = x), corresponding to perfect test-retest agreement.

The test-retest reliability of go errors between the sessions showed good reliability, ICC = .75, 95% CI [.50, .89], and moderate reliability for no-go error rate, ICC = .59, 95% CI [.25, .81].

#### Reaction Time

Figure 4 illustrates RT as a function of session and trial-type, and Figure 5 represents the test-retest reliability between sessions for go and no-go trials. A main effect of trial-type was observed, *F*(1, 22) = 43.45, *p* < .001, *η_p_²* = .66. As expected, go RTs were longer (582 ms) as opposed to no-go stopping time (445 ms), as go trials required movement execution while no-go trials captured the point at which the movement stopped. Similar to error rate neither the session effect, *F*(1, 22) = 2.40, *p* = .135, *η_p_²* = .10, nor the interaction, *F*(1, 22) = 0.09, *p* = .762, *η_p_²* = .004, reached significance.

**Figure 4.**
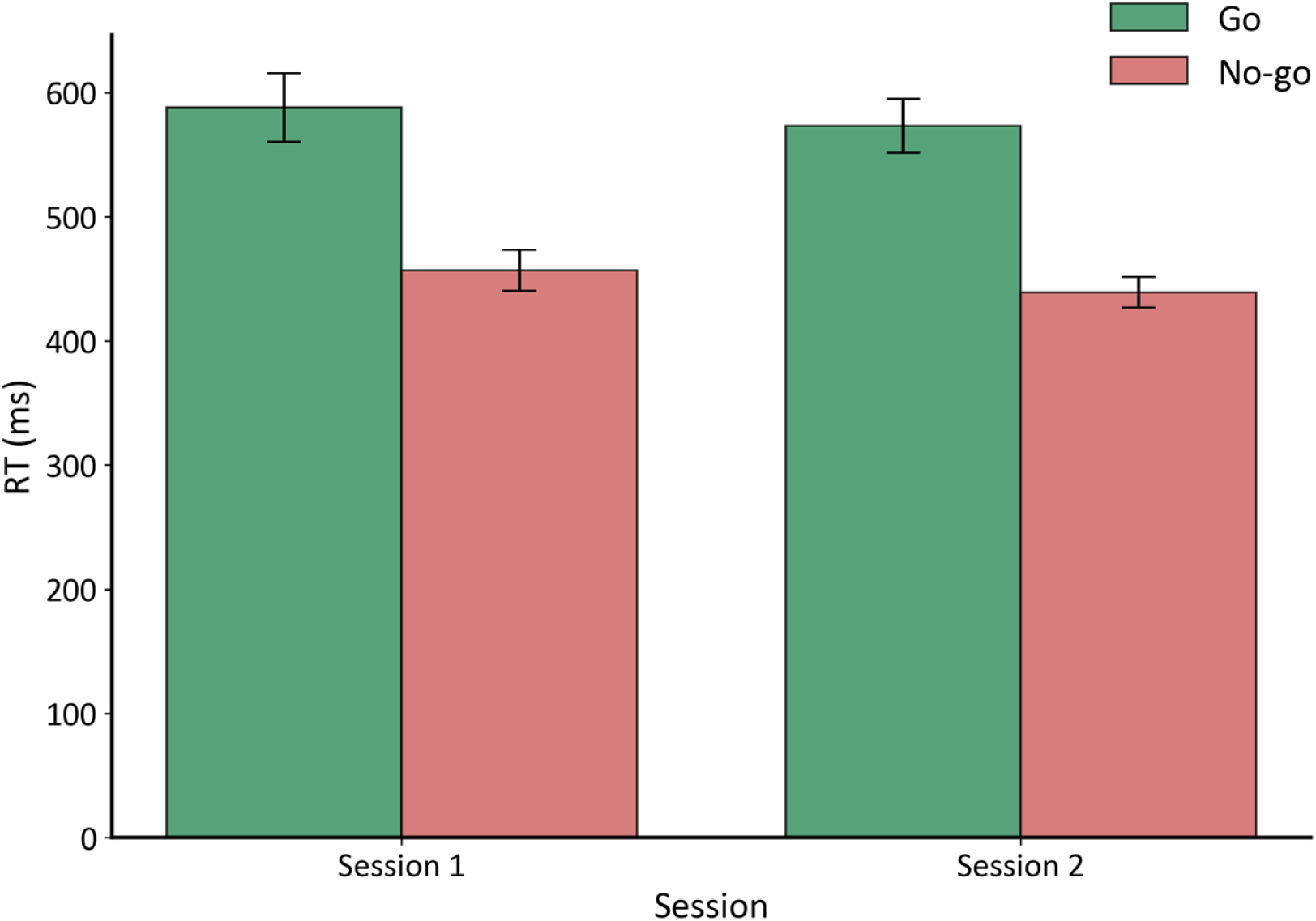
RT by session and trial-type. *Note*. RT (in ms) as a function of session and trial-type. Error bars indicate the standard error of the mean (SEM) across participants.

**Figure 5.**
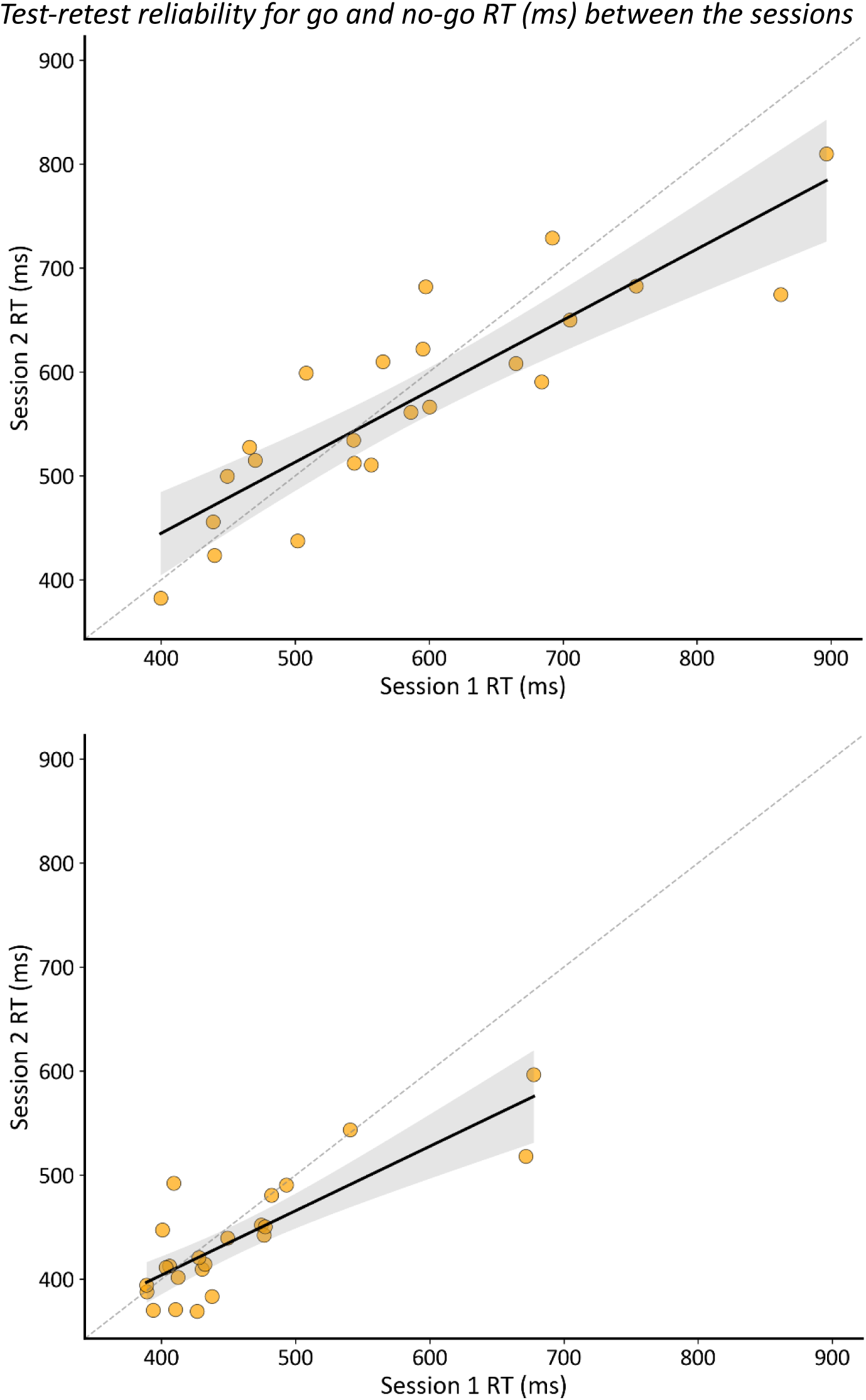
Test-retest reliability for go and no-go RT (ms) between the sessions. *Note.* Scatter plot showing the relationship between session 1 and session 2 RT (ms) for go (top) and no-go (bottom) trials. Each point represents one participant. The black line depicts the ordinary least squares regression of session 2 on session 1, and the shaded region indicates the 95% confidence interval of the mean regression estimate. The dashed diagonal line represents the line of identity (y = x), corresponding to perfect test-retest agreement.

RT showed stronger reliability. Go RT showed ICC = .85, 95% CI [.67, .93], and no-go RT showed ICC = .79, 95% CI [.57, .91], both indicating good test-retest stability.

### 3.2 Kinematics measures

Kinematic analyses were restricted to correct trials only. Outliers were identified and removed using a cell-wise approach (similar to RT) for each kinematic metric, as each metric differs in scale, and extreme observations could be metric-specific. Using this approach, 1.5% of observations were removed for mean velocity, 2.3% for mean acceleration, and 2.4% for path length.

#### Path length

Figure 6 illustrates path length as a function of session and trial-type, and Figure 7 represents the test-retest reliability between sessions for go and no-go trials. A main effect of trial-type was observed, *F*(1, 22) = 32.82, *p* < .001, *η_p_²* = .60. Go trials produced longer path (598 pixels) than no-go trials (542 pixels), reflecting stopping during successful inhibition. Neither the session, *F*(1, 22) = 0.04, *p* = .851, *η_p_²*= .002, nor the interaction, *F*(1, 22) = 0.17, *p* = .682, *η_p_²* = .008, reached significance.

**Figure 6.**
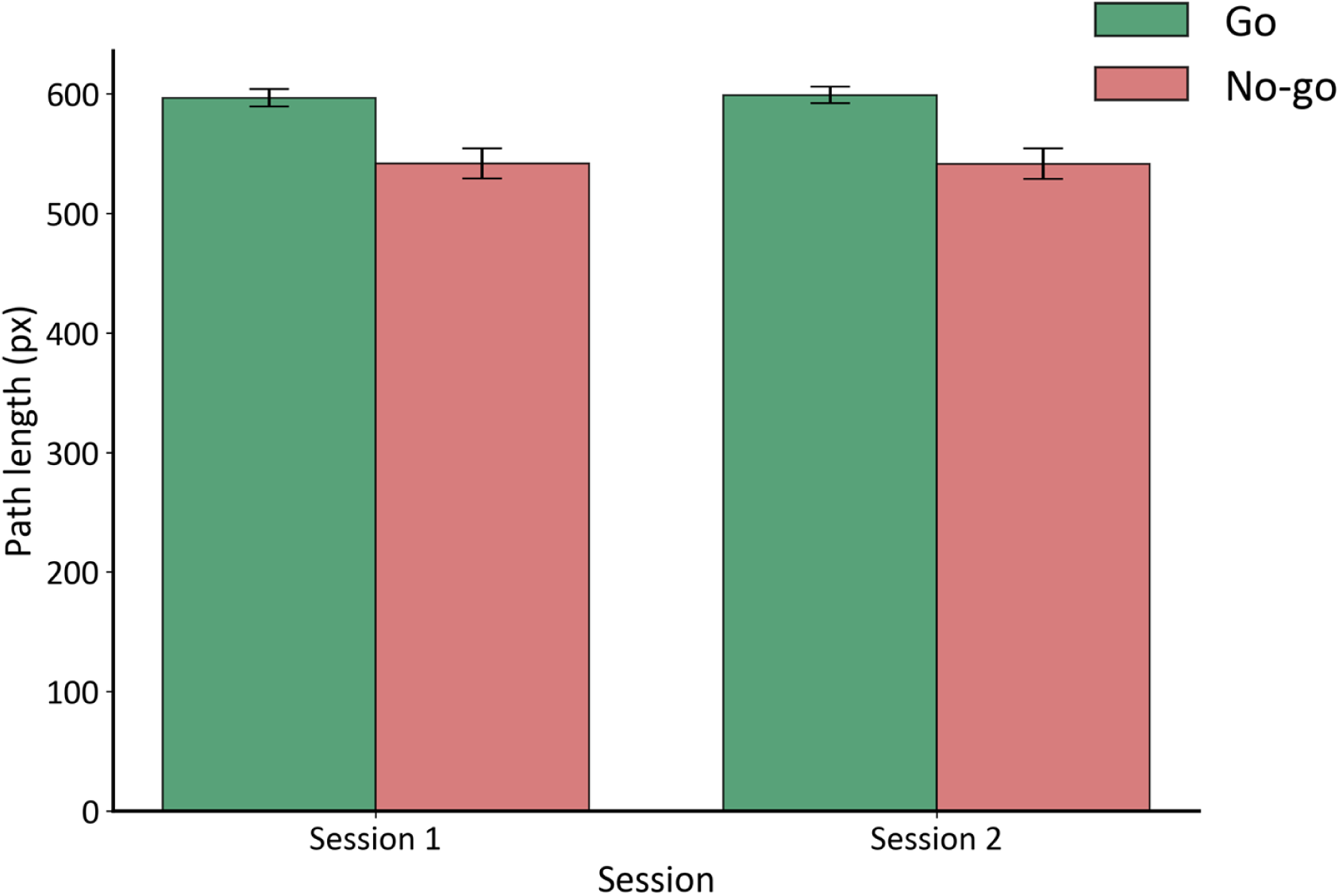
Average Path length by session and trial-type. *Note*. Path length (in pixels) as a function of session and trial-type. Error bars indicate the standard error of the mean (SEM) across participants.

**Figure 7.**
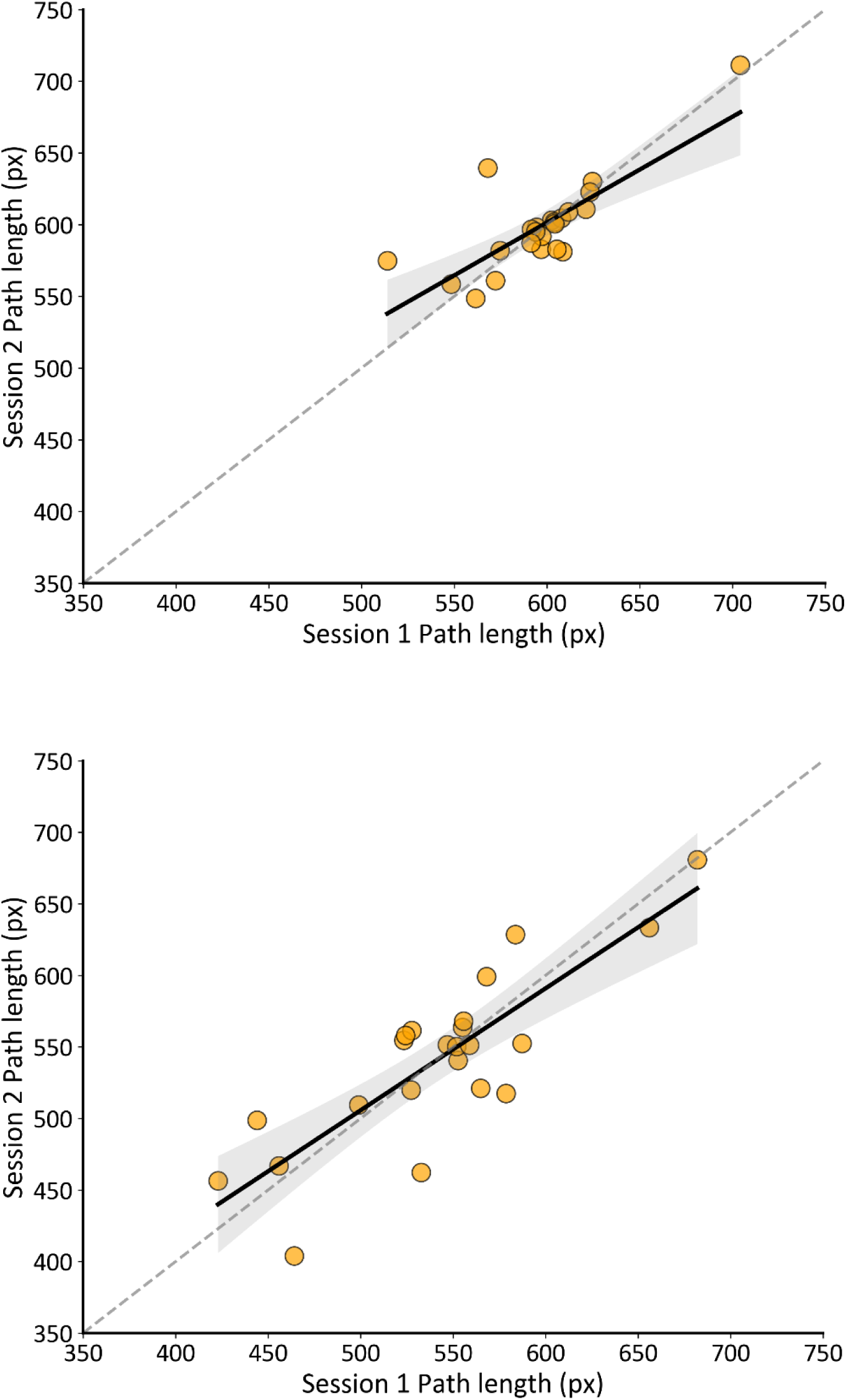
Test-retest reliability for go and no-go path length (px) between the sessions. *Note.* Scatter plot showing the relationship between session 1 and session 2 path length (px) for go (top) and no-go (bottom) trials. Each point represents one participant. The black line depicts the ordinary least squares regression of session 2 on session 1, and the shaded region indicates the 95% confidence interval of the mean regression estimate. The dashed diagonal line represents the line of identity (y = x), corresponding to perfect test-retest agreement.

Path length demonstrated good test-retest reliability for both go trials, ICC = .78, 95% CI [.55, .90], and no-go trials, ICC = .83, 95% CI [.65, .93].

#### Mean velocity

Figure 8 illustrates mean velocity as a function of session and trial-type and Figure 9 represents the test-retest reliability between sessions for go and no-go trials. A main effect of trial-type was observed, *F*(1, 22) = 46.81, *p* < .001, *η_p_²* = .68. Go trials demonstrated higher mean velocity (1091 px/s) compared to no-go trials (934 px/s). Neither the session, *F*(1, 22) = 0.57, *p* = .459, *η_p_²* = .03, nor the interaction, *F*(1, 22) = 0.03, *p* = .854, *η_p_²* = .002, reached significance.

**Figure 8.**
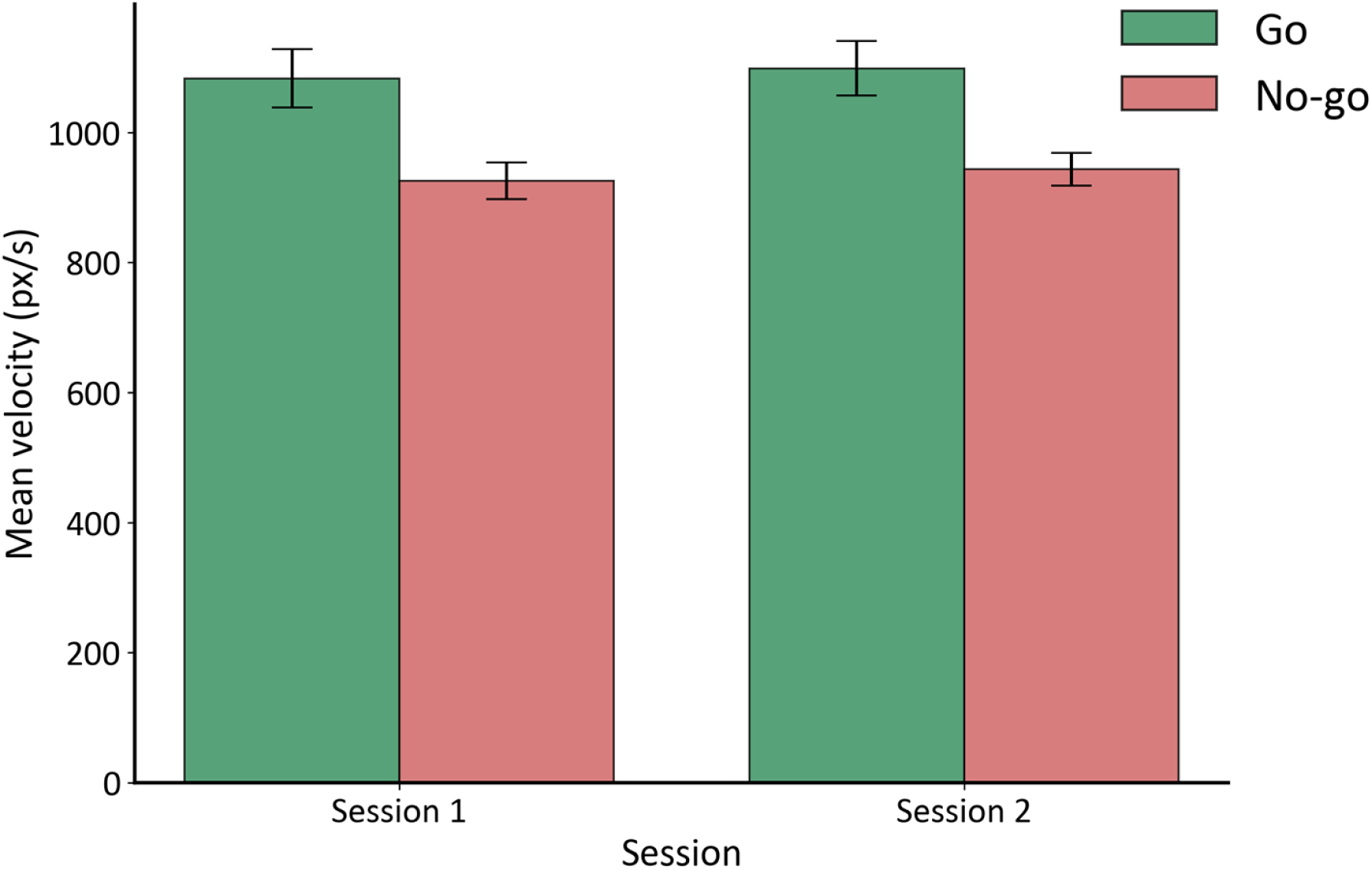
Mean velocity by session and trial-type. *Note*. Bars represent participant-level means computed from correct trials. Error bars indicate the standard error of the mean (SEM) across participants.

**Figure 9.**
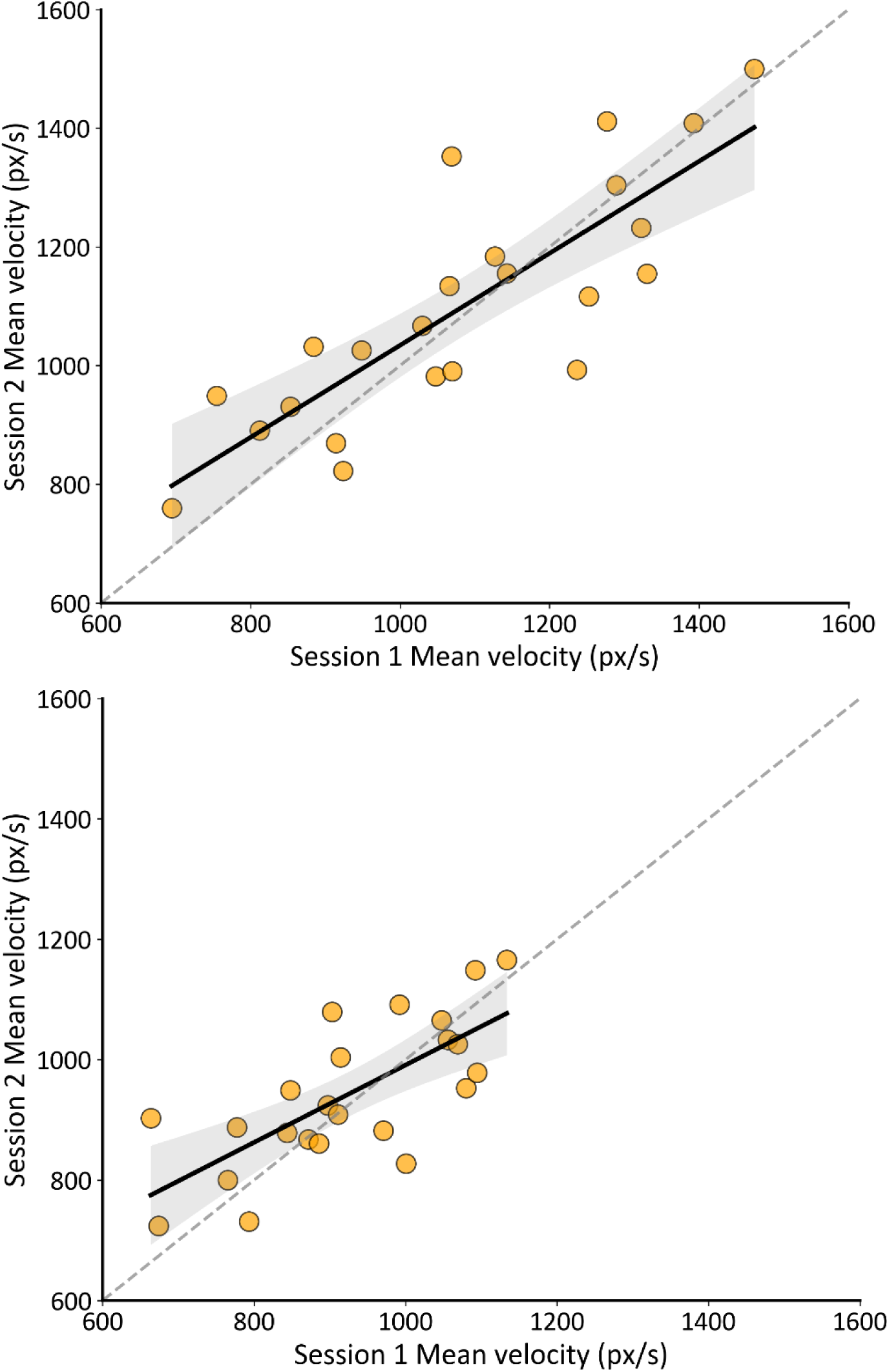
Test-retest reliability for go and no-go mean velocity between the sessions. *Note.* Scatter plot showing the relationship between session 1 and session 2 mean velocity (px/s) for go (top) and no-go (bottom) trials. Each point represents one participant. The black line depicts the ordinary least squares regression of session 2 on session 1, and the shaded region indicates the 95% confidence interval of the mean regression estimate. The dashed diagonal line represents the line of identity (y = x), corresponding to perfect test-retest agreement.

Mean velocity showed good reliability for go trials, ICC = .83, 95% CI [.64, .92], and moderate reliability for no-go trials, ICC = .72, 95% CI [.44, .87].

#### Mean acceleration

Figure 10 illustrates mean acceleration as a function of session and trial-type and Figure 11 represents the test-retest reliability between sessions for go and no-go trials. A main effect of trial-type was observed, *F*(1, 22) = 25.95, *p* < .001, *η_p_²* = .54. Go trials produced higher mean acceleration (25,884 px/s²) than no-go trials (22,264 px/s²), reflecting the greater magnitude of velocity changes required to complete full reaching movements. The session effect was not significant, *F*(1, 22) = 2.81, *p* = .108, *η_p_²* = .11, nor was the interaction, *F*(1, 22) = 0.74, p = .398, *η_p_²* = .03.

**Figure 10.**
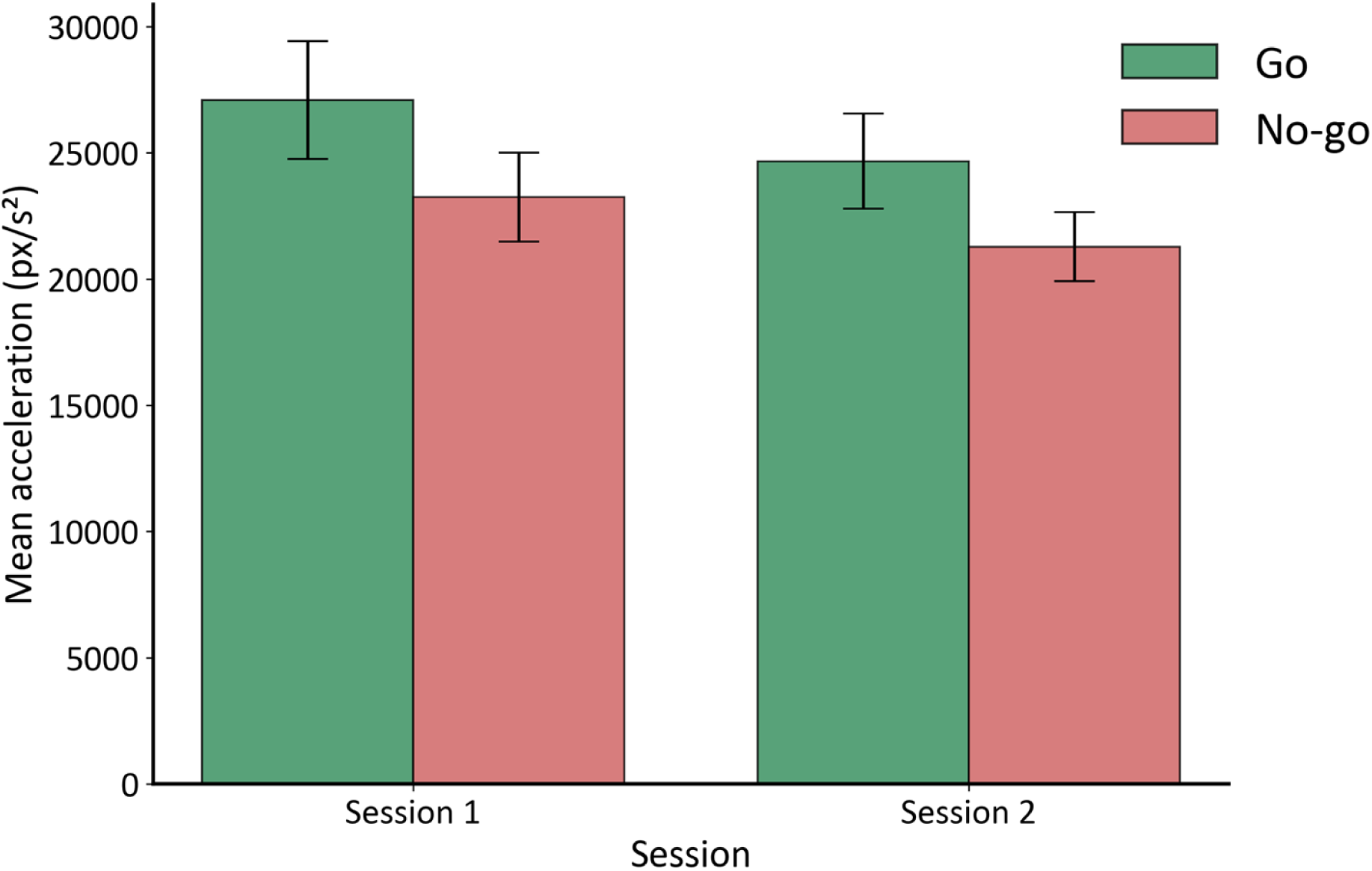
Mean acceleration by session and trial-type. *Note*. Bars represent participant-level means computed from correct trials. Error bars indicate the standard error of the mean (SEM) across participants.

**Figure 11.**
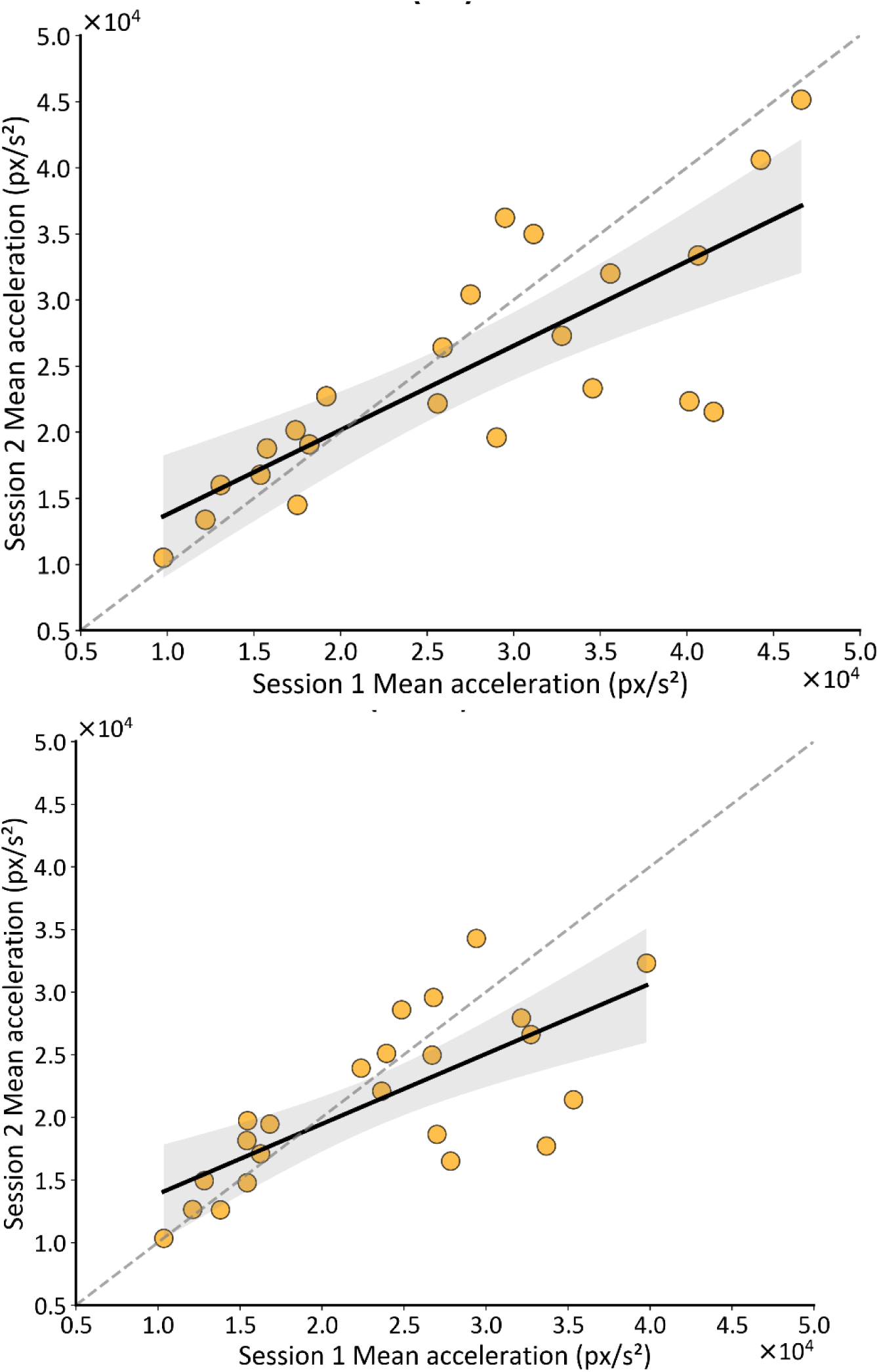
Test-retest reliability for go and no-go mean acceleration between the sessions. *Note.* Scatter plot showing the relationship between session 1 and session 2 mean acceleration (px/s^2^) for go (top) and no-go (bottom) trials. Each point represents one participant. The black line depicts the ordinary least squares regression of session 2 on session 1, and the shaded region indicates the 95% confidence interval of the mean regression estimate. The dashed diagonal line represents the line of identity (y = x), corresponding to perfect test-retest agreement.

Mean acceleration demonstrated good reliability for go trials, ICC = .77, 95% CI [.53, .90] and moderate reliability for no-go trials, ICC = .70, 95% CI [.41, .86].

## 4 General Discussion

The present study developed a mouse-tracking go/no-go paradigm designed to capture inhibitory control during ongoing movement execution. We evaluated both traditional binary measures of inhibitory efficiency (i.e., commission errors) as well as continuous RT and kinematic indices, examining their test-retest reliability across two sessions separated by approximately one week. The results reveal two important findings. First, across all indices, we observed the expected and theoretically sound differences between go and no-go conditions, supporting the construct validity of the paradigm and its capacity to measure inhibition during dynamic motor execution.

Second, all measured indices demonstrated moderate-to-good temporal stability, suggesting the usability of the newly developed paradigm in cross-sectional and longitudinal studies targeting comparable populations. Notably, although the go/no-go paradigm has been shown to yield acceptable reliability for commission errors and go RT (Hedge et al., 2018), the present study demonstrates that continuous kinematic features, particularly path length and RT, show comparable, and in some cases higher, stability (ICCs > .80). Moreover, positive outcomes in young and highly educated individuals encourages future studies validating this paradigm in other cohort, including healthy older cohorts and patient populations with known impairments of inhibitory control.

While traditional go/no-go paradigms typically rely on binary outcomes (i.e., commission errors), the present paradigm provides additional information on continuous RT and kinematic data with demonstrated potential to reveal how inhibition modulates an evolving motor plan. In the current study, we focused on mean velocity, mean acceleration, and path length because they provide an overall summary of how an action unfolds over time and have been widely used in mouse-tracking research to capture response activation and movement dynamics at fine temporal resolution (Kieslich et al., 2019). Mean velocity reflects the overall speed with which a movement is executed, providing a summary of how quickly the motor system commits to the ongoing action.

Mean acceleration captures the rate of change in motor output across the movement epoch, and path length provides a complementary spatial index of how far the movement extended before stopping or completing. Importantly, we observed a clear go/no-go difference across all these kinematic measures. Go trials were characterized by longer path length with significantly greater mean velocity and mean acceleration. In contrast, successful inhibition was characterized by shorter path length and smaller mean velocity and acceleration.

We suggest that this pattern reflects an online modulation of an evolving motor plan (Benedetti et al., 2020). In the go trials, the motor system needs to carry on the movement to reach the target. On the other hand, in no-go trials, inhibitory control is triggered, leading to a rapid reduction in motor drive that brings the movement to a halt. Reductions in velocity and acceleration on no-go trials demonstrate this dynamic recruitment of inhibition. Because our paradigm included substantially more go than no-go trials and required participants to already be in motion before stimulus presentation, participants developed a strong prepotent tendency to respond. This is quite evident in the no-go trials, which was confirmed by the considerable delay that was required before coming to a stop, and participants required measurable time to decelerate to a complete stop.

Nevertheless, participants successfully stopped their movement in no-go trials, tentatively by rapidly modifying their motor plans, demonstrating robust inhibitory control even under these quite challenging conditions. From a motor control perspective, the kinemetric metrics appear to capture distinct aspects of action control. For go trials, the measures likely reflect action commitment, while for no-go trials, these same measures index suppression efficiency. These findings show that our paradigm is capable of systematically capturing online corrections of motor plans to implement response inhibition, which is not always detectable in binary outcomes. Additionally, our results support the notion that inhibition operates as a graded brake on motor vigor rather than an all-or-nothing process (Allain et al., 2004; Cohen & van Gaal, 2014; Ficarella et al., 2019; Neely et al., 2017).

Current neuroscience research has identified the neural pathway that likely mediates the stopping we observed. A recent model proposes that rapid inhibition is implemented through a “hyperdirect” route from frontal cortex to the subthalamic nucleus (STN), supplemented by slower indirect pathways (Jahfari et al., 2010). Critically, research using brain stimulation techniques has demonstrated that the STN is necessary for the broad motor suppression that occurs during stopping, essentially acting as a brake on the entire motor system (Wessel et al., 2022). Our kinematic findings of reduced velocity and acceleration on no-go trials align well with this suppressive mechanism, though these measures alone cannot confirm the specific pathway involved.

Secondly, the present findings show that the measures from this paradigm not only differentiate go and no-go trials but also do so reliably across time. Reliability represents a fundamental psychometric criterion for any cognitive assessment. The go/no-go paradigm has generally shown good reliability (Hedge et al., 2018; Wöstmann et al., 2013). However, the traditional go/no-go paradigm typically assesses the reliability of outcomes such as commission errors and go RTs. The present study extends this work by demonstrating that continuous and kinematic indices also exhibit strong temporal stability, all of which capture theoretically meaningful aspects of inhibitory control. In particular, all measures of go-trial performance showed good reliability (an ICC of between .75 and .85), and no-go measures showed moderate to good reliability. Especially for no-go trials, time to stop (RT) and path length showed good reliability of an ICC of .79 and .83, respectively, while mean velocity, mean acceleration, and error rates showed moderate reliability (ICC of .72, .70, and .59, respectively). The fact that these measures remain stable over time indicates that they index reliable individual differences in inhibitory control. Such stability is a core requirement for longitudinal studies and supports their use in pharmacological or non-invasive brain stimulation research (Wöstmann et al., 2013), that rely on detecting within-subject change over time or in cross-over contexts.

While our findings provide initial evidence for the construct validity and reliability of our paradigm, several methodological considerations need to be discussed. First, the current design included fewer no-go trials than go trials, which, while valid for creating a prepotent response tendency, limits certain statistical analyses, such as sequential modulations of inhibitory control.

However, for the primary goal of developing a reliable paradigm to measure inhibitory control, this design proved effective. Second, in this initial stage of paradigm evaluation, we focused on a selected set of kinematic features, such as mean velocity, mean acceleration, and path length. This was because these features reflect both the strength of action commitment and the extent to which an ongoing movement is suppressed during inhibition. Future research may extend this work by examining additional kinematic features, such as movement curvature or trajectory deviations, to address complementary research questions. Third, the current study established test-retest reliability in a rather small sample of healthy young adults over a one-week interval. Future research should examine whether these reliability estimates replicate in larger sample sizes and generalize to older individuals and clinical populations (e.g., individuals with ADHD or impulse control disorders) and to longer retest intervals, particularly relevant for intervention studies.

In sum, we developed a dynamic go/no-go mouse-tracking paradigm that effectively captures inhibitory control through multiple reliable continuous indices. The observed RT and kinematic differences between go and no-go trials provide compelling evidence for distinct motor signatures of action execution versus suppression. Critically, the moderate-to-good test-retest reliability of these measures establishes their suitability for cross-sectional and longitudinal research in young healthy individuals, with potential for extensions to interventional studies, and the development of behavioral biomarkers of inhibitory control. This methodological advance lays the foundation for future investigations of the temporal dynamics of inhibitory control and applying these sensitive measures in clinical and neuroscience research contexts.

